# High-Fidelity Nanopore Sequencing of Ultra-Short DNA Sequences

**DOI:** 10.1101/552224

**Authors:** Brandon D. Wilson, Michael Eisenstein, H. Tom Soh

**Affiliations:** Department of Chemical Engineering, Stanford University, Stanford, CA 94305, USA; Department of Electrical Engineering, Stanford University, Stanford, CA 94305, USA; Department of Radiology, Stanford University, Stanford, CA 94305, USA; Chan Zuckerberg Biohub, San Francisco, CA 94158, USA

## Abstract

Nanopore sequencing offers a portable and affordable alternative to sequencing-by-synthesis methods but suffers from lower accuracy and cannot sequence ultra-short DNA. This puts applications such as molecular diagnostics based on the analysis of cell-free DNA or single-nucleotide variants (SNV) out of reach. To overcome these limitations, we report a nanopore-based sequencing strategy in which short target sequences are first circularized and then amplified via rolling-circle amplification to produce long stretches of concatemeric repeats. These can be sequenced on the Oxford Nanopore Technology’s (ONT) MinION platform, and the resulting repeat sequences aligned to produce a highly-accurate consensus that reduces the high error-rate present in the individual repeats. Using this approach, we demonstrate for the first time the ability to obtain unbiased and accurate nanopore data for target DNA sequences of < 100 bp. Critically, this approach is sensitive enough to achieve SNV discrimination in mixtures of sequences and even enables quantitative detection of specific variants present at ratios of < 10%. Our method is simple, cost-effective, and only requires well-established processes. It therefore expands the utility of nanopore sequencing for molecular diagnostics and other applications, especially in resource-limited settings.

**One Sentence Summary:** We introduce a simple method of accurately sequencing ultra-short (<100bp) target DNA on a nanopore sequencing platform.

---

Nanopore technology, most notably commercialized as the handheld MinION instrument by Oxford Nanopore Technologies (ONT), has emerged as a powerful sequencing modality due to its low cost, portability, and capacity to sequence very long strands of DNA. These features have made nanopore sequencing indispensable for the assembly of genomes that were previously inaccessible to conventional, short-read, sequencing-by-synthesis methods^1–2^. Despite these positive attributes, nanopore sequencing is ill-suited for important applications, such as the profiling of microRNA^3^ or cell-free DNA^4^, which require sequencing of ultra-short DNA (< 100 bp). In fact, the shortest DNA successfully sequenced on a nanopore platform to date is 434 bp^5,6^. Furthermore, current nanopore sequencing technologies have higher error rates relative to the sequencing-by-synthesis chemistry employed in the widely-used Illumina platforms^7^. Because of this, conventional nanopore sequencing is incompatible with applications that require high accuracy, such as the detection and quantification of single-nucleotide variants (SNVs). To date, the resolution of SNVs on nanopore-based platforms has only been achieved through either very high sequence coverage (>1,000x) or co-sequencing on a high-accuracy, short-read platform such as the Illumina MiSeq^8^. Therefore, a novel methodology that enables sequencing of ultra-short DNA with low error-rates would greatly expand the utility of nanopore sequencing technology.

We describe a simple sample-preparation strategy that converts ultra-short DNA into long stretches of tandem repeats (concatemers) that can be sequenced on the MinION with sufficient accuracy to achieve reliable SNV detection. To the best of our knowledge, this represents the first successful nanopore sequencing of target DNA shorter than 100 bp, as well as the first successful resolution of SNVs from single reads on a nanopore sequencer. Our high-fidelity short reads method (HiFRe) entails the circularization of short DNA sequences, followed by the generation of concatemers via rolling-circle amplification (RCA). These concatemers can be readily sequenced on the MinION, with the tandem repeats providing highly accurate reads through *in silico* reconstruction of the target sequence. Previous work in this area has illustrated that circularizing and concatemerizing DNA is a robust approach to improve the fidelity of sequencing in general^9–11^. Intramolecular-ligated nanopore consensus sequencing (INCseq)^10^, for example, uses blunt-end ligation to circularize double-stranded (ds) DNA of 600-800 bp followed by RCA to create linear DNA with tandem copies of the template molecule. INCseq improves the accuracy of MinION sequencing through computational alignment of the tandem repeats to reconstruct the original sequence. However, its reliance on blunt-end ligation requires the input DNA to be long enough to overcome the curvature-induced strain in double-stranded DNA to achieve circularization, preventing the method from being adapted to ultra-short reads.

HiFRe offers important advantages over this and other previously reported approaches^5,6,10,11^. Most importantly, whereas existing methods for nanopore sequencing of small DNA are still limited to relatively long fragments of 500-1,000 bp, HiFRe enables accurate targeted analysis of sequences < 100 bp. As a demonstration, we have used HiFRe to sequence targeted 52-bp segments of DNA with high fidelity. Furthermore, we show that HiFRe enables SNV resolution from a single nanopore read, with the capacity to accurately quantify mixtures of sequences based on the discrimination of single-nucleotide differences at ratios as low as 10:90.

## Results and Discussion

HiFRe enables sequencing analysis of ultra-short DNA targets through a straightforward procedure of circularization and amplification (**Figure 1a**). First, we employ molecular inversion probes (MIPs)^12,13^ to copy a target DNA sequence into a circular single-stranded (ss) DNA construct. For this reaction, we employ the Phusion polymerase (New England Biolabs), which lacks 5’**→**3’ exonuclease activity, thereby ensuring that extension halts at the 5’ end of the second hybridization site, where ligation is to occur. We also took care to design MIPs that are predicted to exhibit minimal secondary structure—especially in the anchor sites—in order to ensure highly efficient circularization. We established a threshold for secondary structure energetics based on the Δ*G*_*folding*_ for which the conversion from linear to circular DNA is >90% after five temperature cycles. In order to achieve efficient circularization, we estimate that any secondary structure involving an anchor site must have Δ*G*_*folding*_ ≥ −0.33 *kcal*/*mol* (**Supplemental Calculation 1**). Prior works that did not consider this aspect of MIP design used 99 cycles of melting, annealing, extension, and ligation to achieve circularization^13^, whereas we are consistently able to obtain complete conversion to circular product in just five cycles (**Figure 2**). After ligation, we incubate the reaction with a mixture of exonucleases I and III to degrade any remaining linear single- or double-stranded DNA while leaving circular DNA intact. We note that this combination of circularization and degradation results in excellent specificity, allowing us to efficiently isolate and purify target amplicons even in a large background of non-specific byproducts of PCR amplification (**Supplemental Figure 1**).

**Figure 1:**
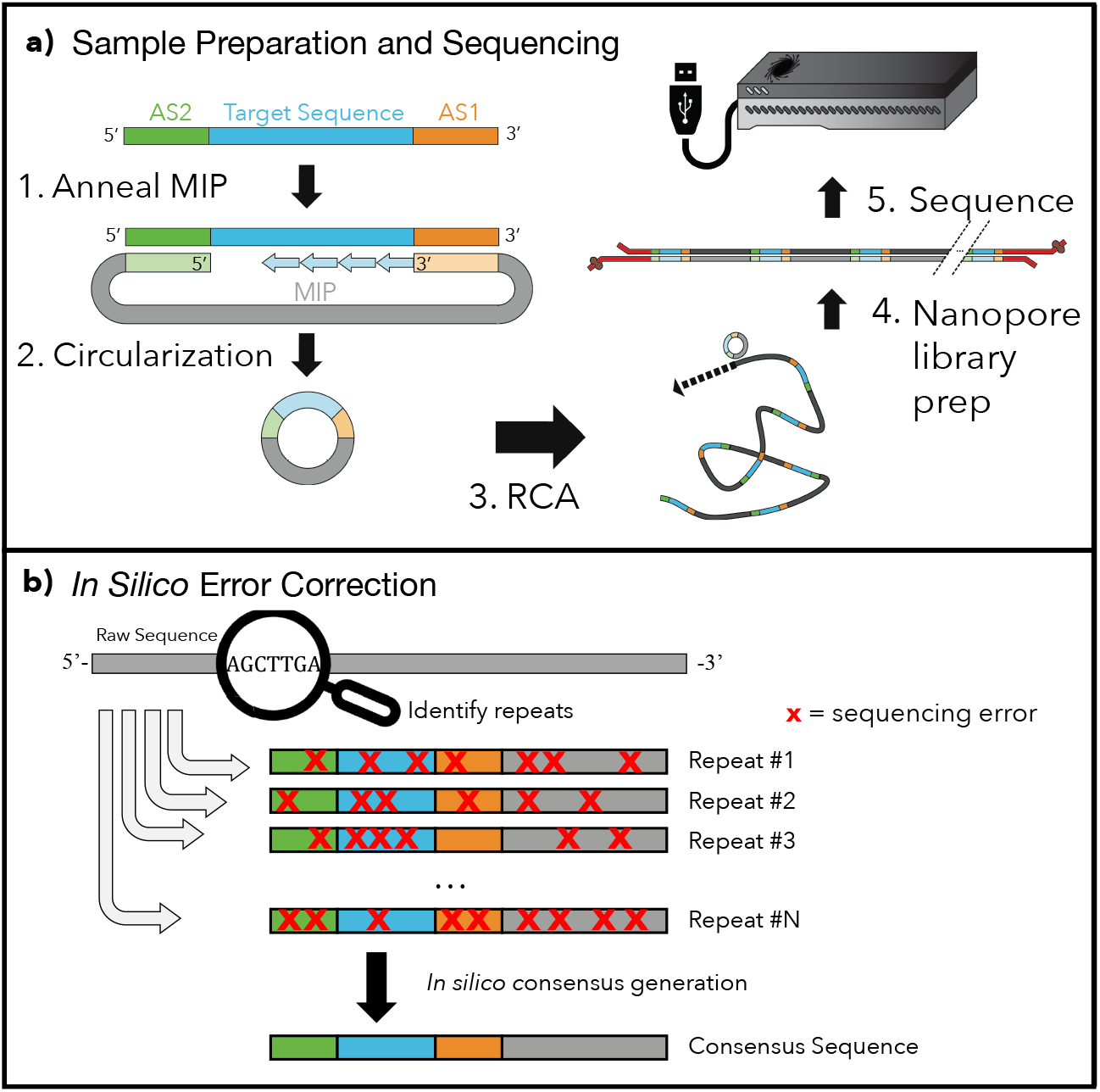
Sequencing ultra-short reads on the MinION. **a)** 1) Molecular inversion probes (MIPs) are annealed to the target sequence (blue) at anchor site 1 (AS1, orange) and anchor site 2 (AS2, green). Phusion polymerase copies the target sequence into the MIP; the lack of 5’**→**3’ exonuclease activity ensures that extension halts when the polymerase reaches AS2. 2) Ampligase ligates the extended template to the phosphorylated 5’ end of the MIP, generating circular ssDNA. Linear ss- or dsDNA fragments are degraded by a combination of exonuclease I and exonuclease III. 3) The circular DNA is subjected to RCA to generate tandem repeats of the original target, yielding ultra-long, concatemerized ssDNA. 4) The RCA product is converted to dsDNA with Taq polymerase and subjected to ONT library preparation. 5) Sequencing reads are collected from a new MinION R9.4 flow-cell run for 24 hrs. **b)** The raw sequences are compiled and analyzed. The identified repeats have poor accuracy in isolation, but since the sequencing errors vary across repeats, they can be aligned together to produce a high-fidelity consensus sequence.

**Figure 2:**
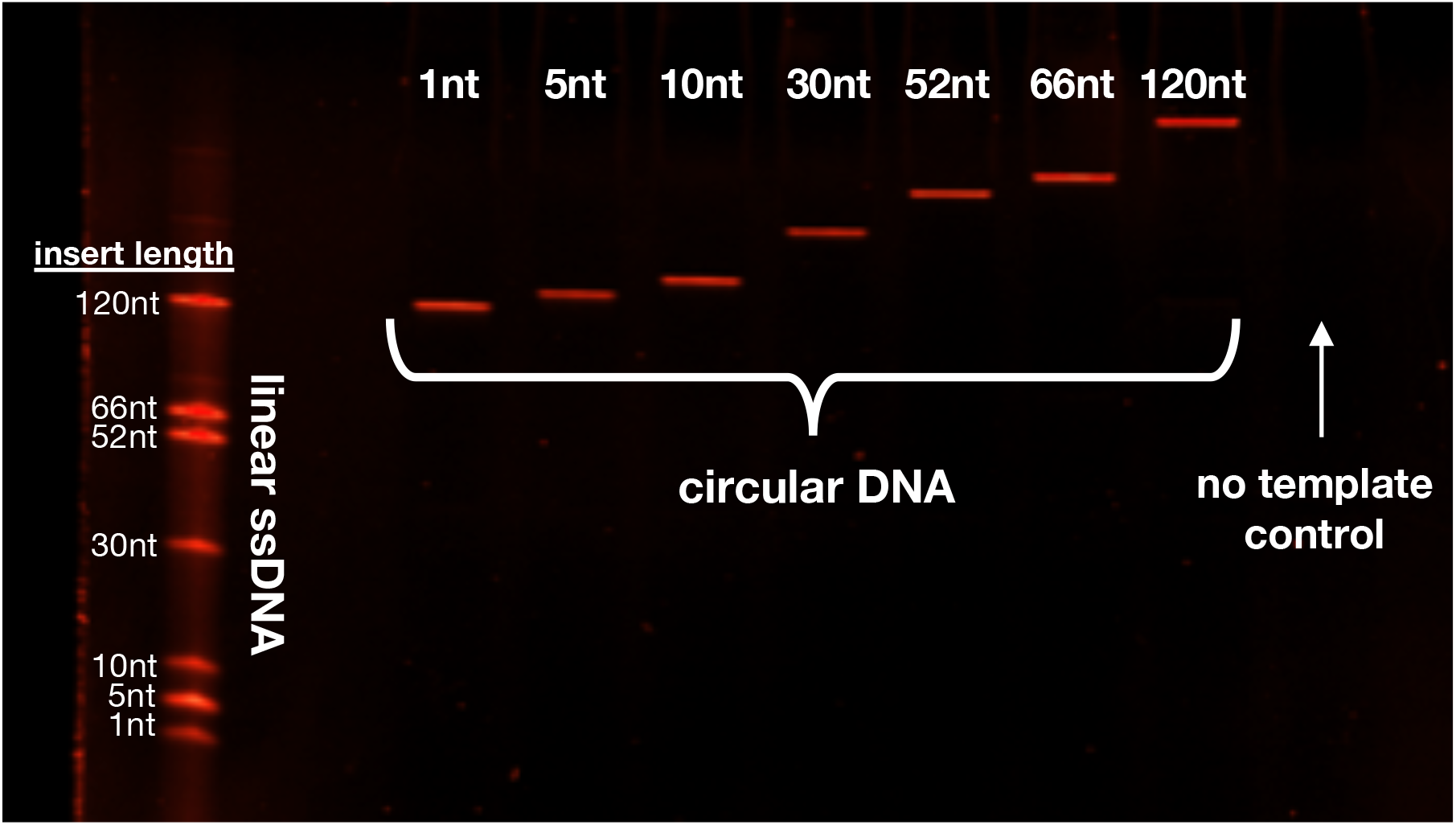
Circularization efficiency is independent of target sequence length. After five rounds of temperature cycling and subsequent exonuclease treatment, we achieve consistently efficient circularization for target sequences ranging in length from 1 to 120 nt (lanes 3–9). Lane 1 contains a mixture of all of the linear ssDNA target sequences. The lengths listed are the lengths of the target region; the full lengths in lane 1 have additional flanking 28- and 23-nt anchor sites, and the full lengths in lanes 3-9 have an additional 102 nt from the MIP itself.

We then initiate RCA^14^ on the circular product with the strand-displacing polymerase *Φ*29, which generates long single-stranded concatemers (>1,000 kb) of the complement to the circular template. Shorter template circles are generally advantageous here, as they can generate more repeats within a fixed RCA length, which in turn positively impacts the accuracy of the resulting computational alignment. There are certain limitations, including the fact that the ring strain in very small DNA circles can cause *Φ*29 to dissociate prematurely^15^. However, since the MIP itself is 102 nt, HiFRe circumvents this limitation and can target inserts as short as a single nucleotide. The ssDNA generated by the RCA reaction is incompatible with the dsDNA required by the MinION library preparation protocol. We therefore implement a brief secondary amplification with Taq polymerase to generate the required starting material of ∼1 µg dsDNA. The presence of multiple primer sequences within the concatamer creates opportunities for laddering and shortening in the amplification product pool. We exploit the 5’**→**3’ exonuclease capacity of Taq polymerase to overcome this problem: any sequences that are undesirably primed from an internal primer-binding site are likely to be degraded by the exonucleolytic activity of an upstream Taq enzyme, greatly favoring the synthesis of full-length products. Finally, the dsDNA products are prepared for sequencing via ONT’s standard 1D amplicon by ligation kit and sequenced on a R9.4 flow cell. The tandem repeats identified from the raw reads are then used to computationally reconstruct the original sequence (**Figure 1b**).

### HiFRe enables accurate MinION sequencing of ultra-short DNA sequences (< 100 bp)

To demonstrate the generalizability of this approach, we first performed our circularization procedure on a variety of sequences with lengths ranging from 1–120 bp. We found that all of these can be targeted and circularized with high efficiency, using the same 102-bp MIP design for each target. The observation that high circularization efficiency is preserved independent of the tested target length (**Figure 2**) illustrates the generalizability of this approach over a range of ultra-short sequence lengths.

In order to further explore the performance and capabilities of our HiFRe method, we subjected a 52-bp target sequence (see **Table 1** for this and other sequences used in this work) to the entire workflow, including sequencing and data analysis. We circularized our target sequence with a 102-nt MIP, yielding an RCA product with a repeat length of 154 bp. To account for potential sequence-induced biases that could arise from using a single test sequence, we incorporated two randomized N_10_ regions at the ends of the 52-bp target sequence, adjacent to the anchor sites. Notably, the rest of the workflow can be adapted to any other assay that also results in a circular DNA output, such as MiR-ID^16^ or cPLA^17^.

**Table 1:**
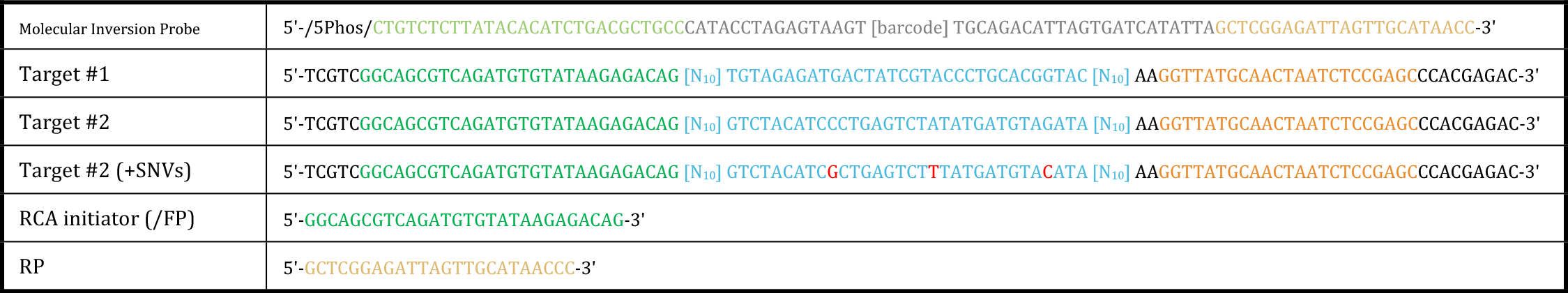
DNA sequences used in this work. Sequences in blue represent the target sequences, green and orange represent universal anchor sites 1 & 2, respectively, and grey represents the MIP scaffold sequence. Bases in red indicate the three SNVs that were introduced into Target #2. The barcode in the MIP sequences is a 12 nt region that allows for multiplexing in a single nanopore Barcode sequences are listed in Supplementary Table 1

We subsequently demonstrate that the resulting consensus reads achieve excellent accuracy despite the short length of the original template. In general, short reads are less tolerant of errors than long sequences in terms of sequence identity. For example, an *E. coli* genome sequence with a 20% error rate will be accurately identified as *E. coli* since there are still millions of accurately-sequenced bases and the sequence space is very sparse. On the other hand, a 20-bp miRNA with a 20% error rate has a high probability of being misidentified as a completely different miRNA, as the sequence space is far more dense^18^. Therefore, we developed a computational algorithm to process the data from HiFRe, identify tandem repeats in the raw base-called reads, and reconstruct the original sequence. The source code for this algorithm is publically available on GitHub (see Methods).

A representative alignment illustrates the utility of using individual tandem repeats to reconstruct an accurate consensus sequence (**Figure 3a**). In order to recover the individual repeats, the algorithm scans the raw base-called reads for regions that map to the RCA initiator. These individual repeats are compared to the initial target sequence in order to define the average unmapped accuracy. Using a progressive multiple alignment algorithm, we then generate a consensus sequence from the ensemble of consecutive repeats. The consensus sequence is compared to the original target sequence to define the post-alignment accuracy. We observed marked improvement in the alignment scores after consensus sequence generation, with the median alignment score increasing from 0.65 to 0.87 after alignment (**Figure 3b**). It should be noted that we applied the algorithm to *any* read that had more than three identifiable repeats, regardless of the quality score or pass/fail designation reported by the MinKNOW software (in stark contrast to previous methods that only consider high-quality, passing reads^10^).

**Figure 3:**
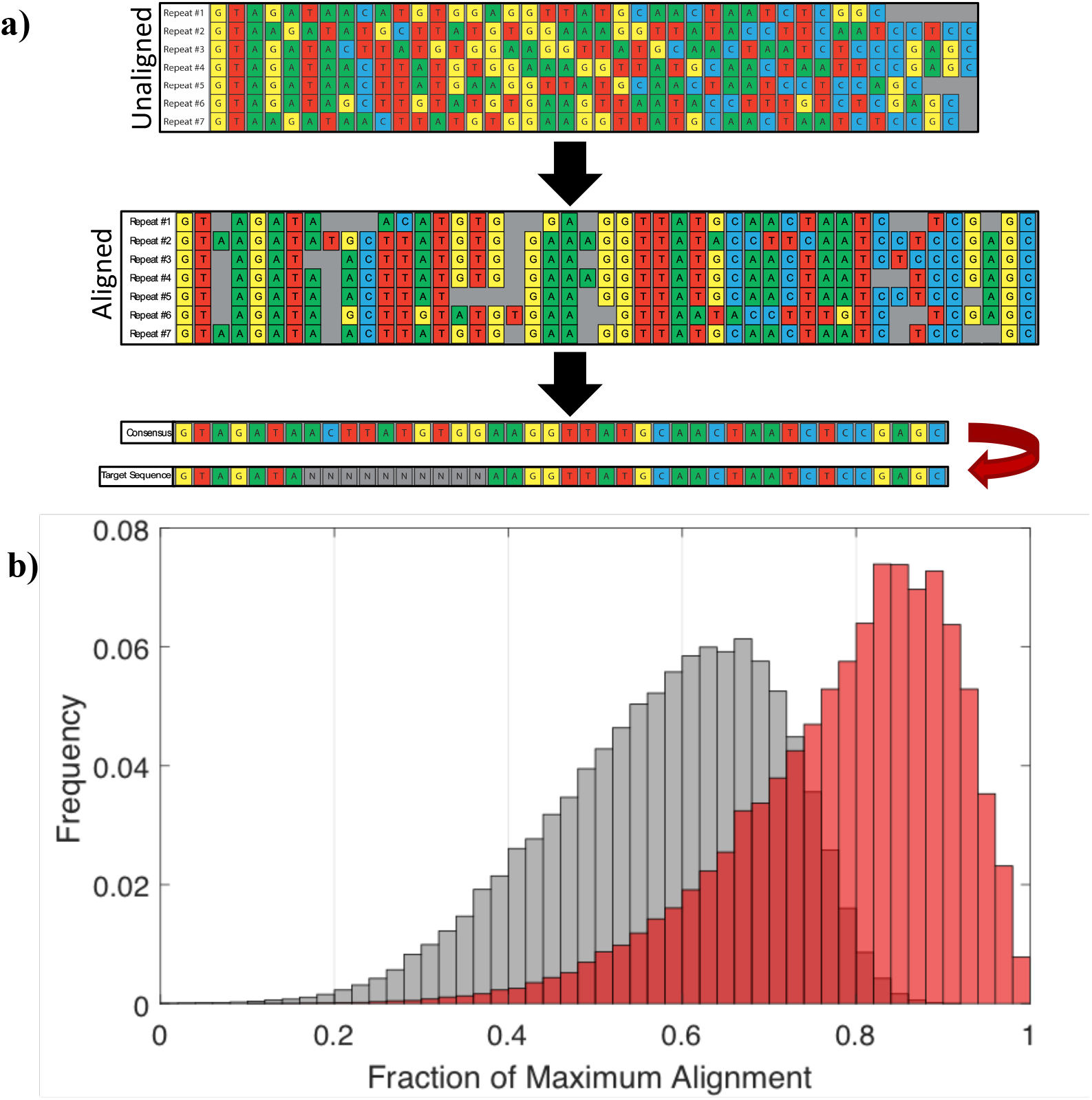
Improving read accuracy through repeat-based consensus. **a)** Representative consensus sequence generation. A single, base-called read is split into its individual repeats. These repeats are aligned with each other to generate a consensus sequence via a winner-take-all base-calling strategy. The consensus sequence is then compared back to the original sequence to assess post-alignment accuracy. **b)** Histogram of alignment scores before (grey) and after (red) consensus sequence generation. The “before” alignment score is an average over the alignment scores of all the repeats found within a single raw read. Data includes all reads with more than three identified repeats, regardless of the quality score or pass/fail designation of the MinION software.

As expected, we found that the improvement in alignment is a strong function of the number of tandem repeats used to generate the consensus sequence (**Figure 4**). Contrary to previous reports that show a plateau in accuracy at 15 repeats^10^, we found that we could achieve significant increases in accuracy with up to 30 repeats. Further gains may be possible beyond that, but the read depth at these lengths was too low to determine the statistical significance of any further increases in accuracy. Improvements to the RCA protocol^19^ could enable the production of higher fractions of longer concatemers, and therefore yield an even more pronounced increase in overall accuracy. Additionally, we observed that the standard deviation of the alignment score also decreases with greater numbers of repeats (**Figure 4**), yielding alignments that are both more accurate and more precise.

**Figure 4:**
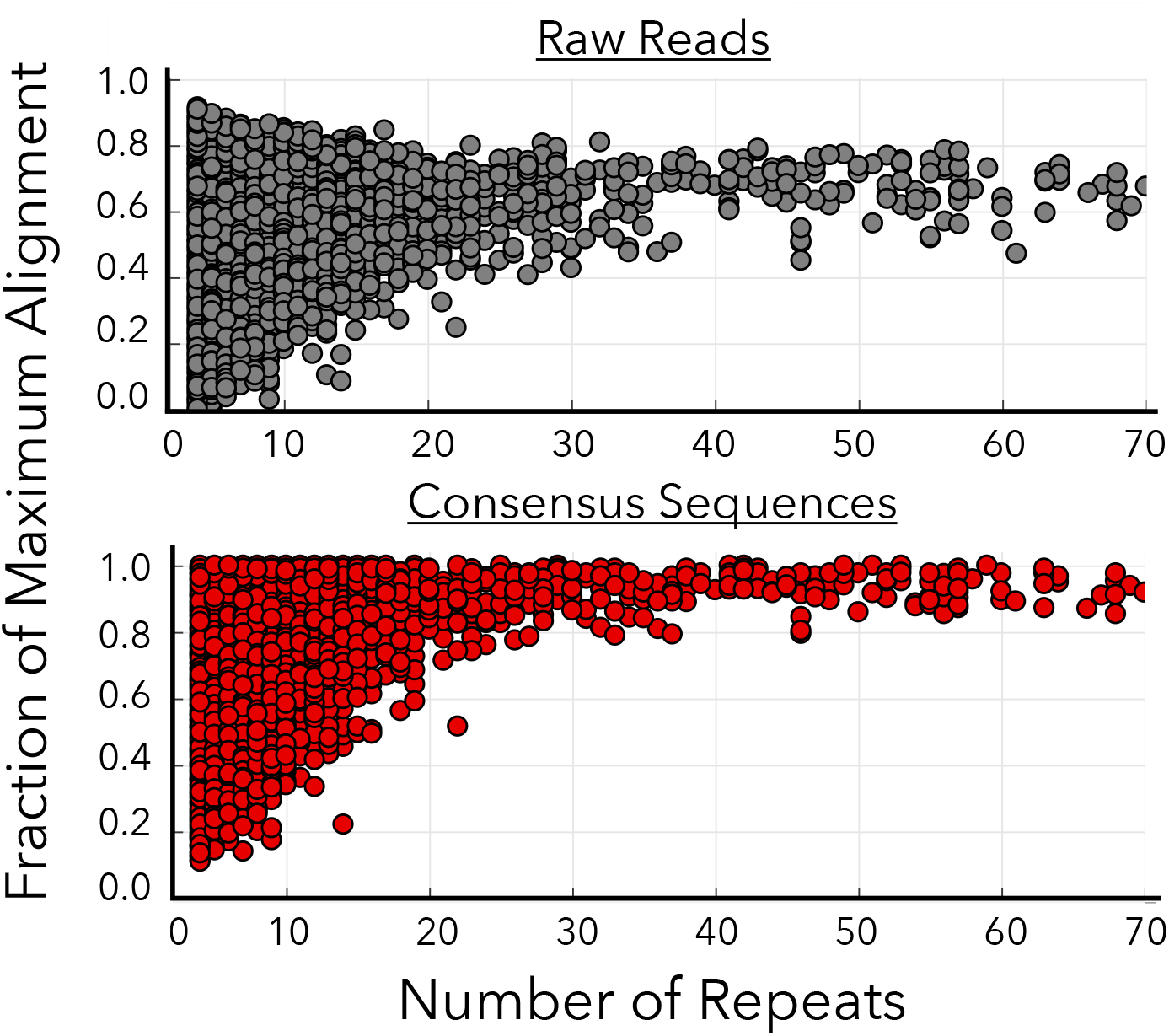
Increased accuracy from alignment of tandem repeats. Plots show normalized Smith-Waterman alignment scores as a function of the number of repeats before (grey) and after (red) alignment. Before consensus sequence generation, alignment score exhibits no dependence on repeat count. Since each “before point” represents an average over all repeats in that read, the observed narrowing arises solely because the increased number of repeats decreases the standard deviation of the average alignment score. After the consensus sequence is generated, the alignment accuracy exhibits a strong dependence on the number of repeats used.

### HiFRe enables quantitative discrimination of sequence variants

Our method is also quantitative, enabling ratiometric quantification of different target sequences. This is useful for many sequencing-based assays, such as transcriptome analysis^20^, miRNA profiling^21,22^, and cfDNA detection^4^. Most uses of nanopore sequencing to date have generally not explored molecular counting, except for quantifying chromosome copy number for aneuploidy detection^5^. RNA-seq can be performed with the nanopore system^23–24^, but this application is still in the nascent stages of commercial development. We have found that HiFRe can accurately quantify input ratios of two 52-nt target sequences ranging from 0 to 90% with an R^2^ of 0.993 (**Figure 5a**), yielding a limit of detection^25^ of 3.3 ± 2.1%. This strong linear relationship between input and output abundances illustrates the power of HiFRe for quantifying relative abundances of multiple sequences.

**Figure 5.**
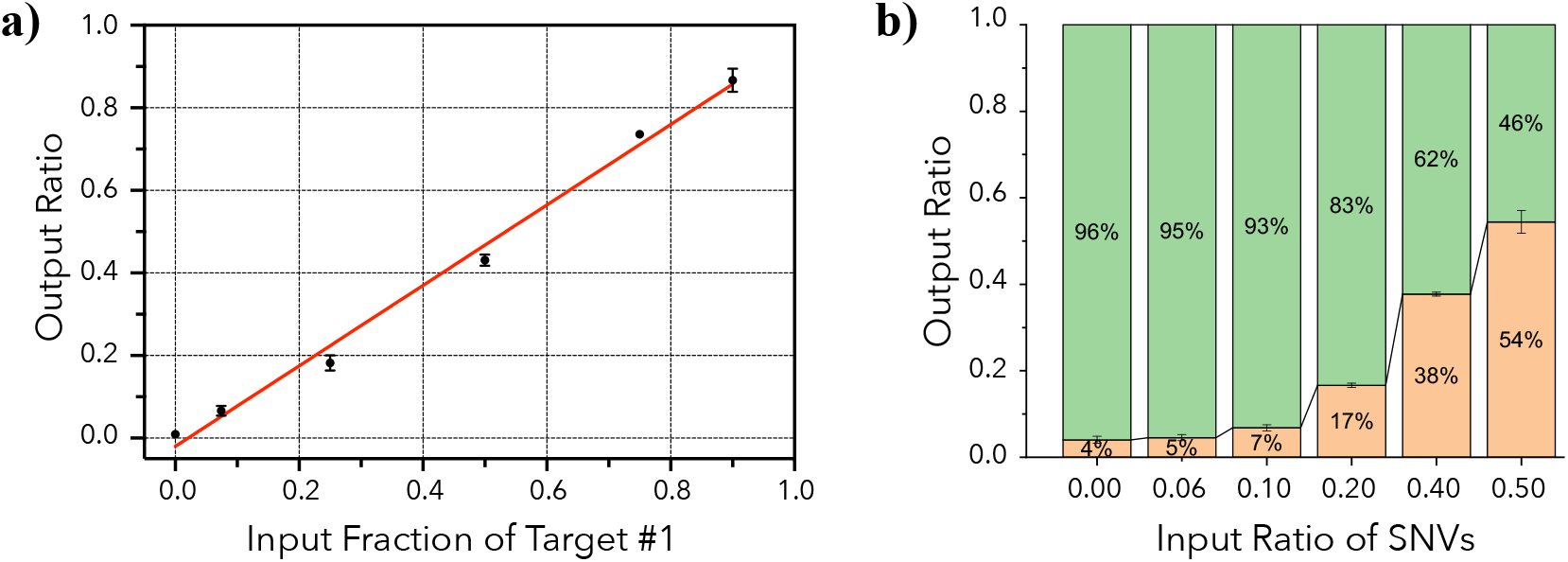
Quantitative analysis with HiFRe. a) Counting relative molecular abundance for two sequences present in mixtures at different ratios. A linear fit to *y* = *mx* + *b* yielded *m* = 0.97 ± 0.04 and *b* = −0.02 ± 0.02 with *R*^2^ = 0.993. This strong linear relationship results in a limit of detection^25^ of 3.3 ± 2.1%. b) Discrimination and quantitation of SNVs in short DNA sequences. Two sequences differing by three SNVs were mixed together in different ratios, and the plot shows the output ratios recovered after HiFRe analysis. Green bars represent the fraction of the original sequence and yellow bars the sequence with three SNVs. In both panels, the error bars represent the standard deviation in the mean of two multiplexed sequencing runs.

This discriminatory power is important, as many clinical applications of DNA sequencing involve resolving the relative abundance of sequences with single-nucleotide differences—for example, the detection of cancerous mutations in cell-free DNA^4^ or the analysis of closely-related miRNA families^21,22^. Until recently, SNV resolution on nanopore-based platforms has required either extremely high read depth or co-sequencing with another sequencing method. HiFRe now enables the resolution of SNVs from single reads within a standard MinION experimental workflow. To demonstrate this, we mixed together two 52-bp test sequences that were identical apart from three nucleotides. Using HiFRe, we were able to accurately resolve the representation of the two sequences for all three SNVs at a variety of different input ratios (**Figure 5b**). Although we tested three different SNVs (G**→**C, A**→**T, C**→**G) to account for potential biases in nanopore base-calling errors^26^, we observed no impact on sequence discrimination from these individual SNVs.

## Conclusion

HiFRe offers a simple and straightforward strategy for the accurate nanopore sequencing of ultra-short sequences (<100 bp). This method deploys RCA on a MIP-circularized template in order to generate long concatemers that can readily be sequenced on the MinION platform. Each raw read contains numerous repeats of the original target sequence that can be computationally aligned together to generate a highly accurate consensus sequence. Using a 52-bp target sequence, we demonstrate over multiple experiments that the accuracy of this method is sufficiently high to enable relative molecular counting and SNV resolution, resulting in accurate ratiometric quantification of DNA sequences that are present at input ratios below 10%. With the added ability to analyze short targets, nanopore sequencing can be used not only for genomic analysis but as a readout for the many bioassays that feature the production of short DNA oligonucleotides as an output, such as PLA^27^, PLAYR^28^, and AbSeq^29^. Ultimately, assays that previously required a fully equipped laboratory and a desktop sequencer could potentially be performed at a fraction of the cost in a portable format, greatly expanding their utility and accessibility, especially in resource-limited settings.

## Methods

### Reagents

Unless noted otherwise, all DNA was purchased from Integrated DNA Technologies and all reagents were purchased from New England Biolabs. Initial DNA concentrations were measured and normalized to 100 μM via A_260_ measurement with a Nanodrop 2000 (Thermo Scientific).

### Probe design

Probes were carefully designed to minimize secondary structure in the hybridization regions using a combination of the NUPACK^30^ and *mfold*^31^ DNA folding applications. We established a threshold of >90% conversion from linear to circular DNA after five cycles of annealing, extension, and ligation. As described in **Supplementary Calculation 1**, this means that any secondary structures in the MIP’s anchor sites must have a Δ*G* > −0.33 kcal/mol. This was an important consideration for the MIP design as a whole as well as all the barcodes listed in **Supplementary Table 1**.

### Preamplification/generation of dsDNA input templates

dsDNA was generated from the ssDNA mixtures listed in **Table 2**. Standard PCR reactions were prepared as follows: 18 μL ssDNA input, 60 μL 2x GoTaq Master Mix (Promega), 12 μL 10 μM forward primer (FP), 12 μL 10 μM reverse primer (RP), and 18 μL nuclease-free water (Ambion). Reactions were carried out by thermocycling with an initial denaturation at 95 °C for 3 min and 20 cycles of 95 °C for 10s, 55 °C for 30s, and 72 °C for 30s on a vapo.protect Mastercycler Pro thermocycler (Eppendorf). After clean-up with a MinElute PCR Purification Kit (Qiagen) via the manufacture’s protocol, amplified dsDNA product was normalized to 100 nM in nuclease-free water. The resultant amplified dsDNA simulates a typical input to the HiFRe workflow (**Supplemental Figure 1**, top).

**Table 2:**
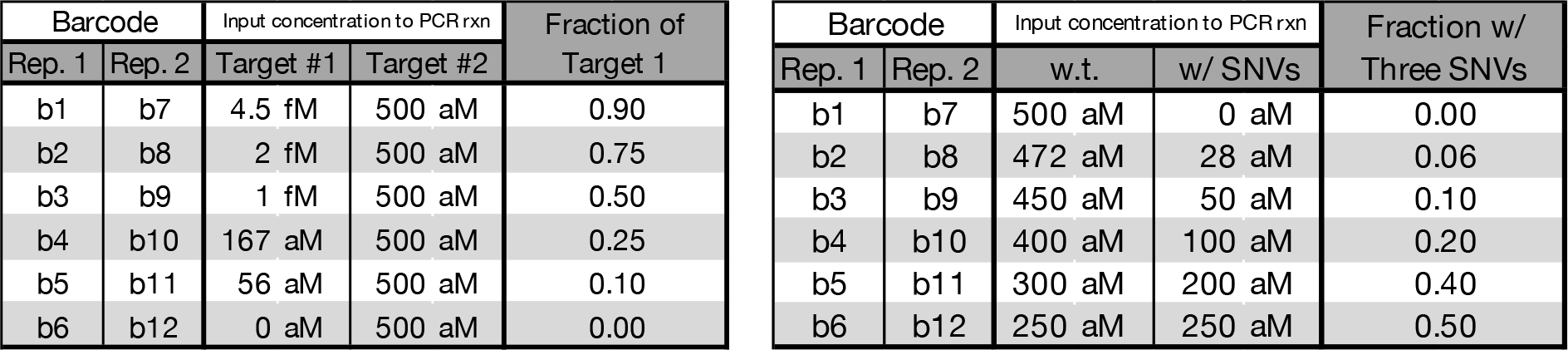
Experimental conditions for a) ratiometric and b) SNV experiments. The barcode represents the sequence used in the N_12_ region to demultiplex the pooled samples (**Supplementary Table 1**). Values shown represent the concentrations in the mixture added to the PCR reaction.

Experimental Conditions:
| Barcode |  | Input concentration to PCR rxn |  | Fraction of Target 1 |
| --- | --- | --- | --- | --- |
| Rep. 1 | Rep. 2 | Target #1 | Target #2 |  |
| b1 | b7 | 4.5 fM | 500 aM | 0.90 |
| b2 | b8 | 2 fM | 500 aM | 0.75 |
| b3 | b9 | 1 fM | 500 aM | 0.50 |
| b4 | b10 | 167 aM | 500 aM | 0.25 |
| b5 | b11 | 56 aM | 500 aM | 0.10 |
| b6 | b12 | 0 aM | 500 aM | 0.00 |

| Barcode |  | Input concentration to PCR rxn |  | Fraction w/ Three SNVs |
| --- | --- | --- | --- | --- |
| Rep. 1 | Rep. 2 | w.t. | w/ SNVs |  |
| b1 | b7 | 500 aM | 0 aM | 0.00 |
| b2 | b8 | 472 aM | 28 aM | 0.06 |
| b3 | b9 | 450 aM | 50 aM | 0.10 |
| b4 | b10 | 400 aM | 100 aM | 0.20 |
| b5 | b11 | 300 aM | 200 aM | 0.40 |
| b6 | b12 | 250 aM | 250 aM | 0.50 |

### Circularization

Optimal MIP annealing temperature was determined to be 60.4 °C based on yield after circularization with a gradient of annealing temperatures ranging from 58–70 °C on a CFX96 Real-Time qPCR System (Bio-Rad). For all of these experiments, 15 nM target dsDNA was circularized with 20 nM MIP. 10 μL circularization reactions were prepared on ice containing 1.5 μL 100 nM target, 2 μL 100 nM MIP, 5 μL 2X Phusion Master Mix, 0.5 μL ampligase (10 U/μL, Lucigen), and 1 μL 10X ampligase buffer (Lucigen). Circularization was achieved under the following conditions: 95 °C for 3 minutes followed by 5 cycles of 95 °C for 30s, 60.4 °C for 1 min, and 37 °C for 2 min. After circularization, linear DNA was degraded by adding 0.45 μL of exonuclease I (20 U/μL) and 0.45 μL of exonuclease III (100 U/μL). The reaction was incubated at 37 °C for 90 min followed by heat inactivation at 65 °C for 20 min. Exonuclease inactivation was determined to be sufficiently efficient based on the lack of exonuclease activity observed in a sample left at 37 °C for 2 days, thus no clean-up step was performed prior to the RCA reaction.

### Rolling circle amplification

The RCA reaction was performed under the following conditions: 2 μL nuclease-treated circularization product, 1 μL *Φ*29 (10 U/μL), 1 μL 10 μM RCA FP, 4 μL dNTP mix (2 mM each, Novagen), 2 μL 10x *Φ*29 buffer, 7.8 μL nuclease-free water, and 0.2 μL 100X BSA. Reactions were carried out with 12 hr incubation at 30 °C, 10 min at 60 °C, and a hold at 4 °C. We also tested 2-, 4-, and 12-hr RCA reactions, and observed no difference in median product length, which indicates that the protocol could be shortened substantially with further optimization of the RCA protocol.

We then amplified the single-stranded RCA product to dsDNA with standard PCR using the following reaction conditions: 9 μL RCA product, 15 μL 2x GoTaq Master Mix (Promega), 1.5 μL 10 μM FP, 1.5 μL 10 μM RP, and 3 μL nuclease-free water. The reaction was initiated with a 2 min hot start (95 °C) followed by 5–8 cycles of 95 °C for 10 s, 58 °C for 30 s, and 72 °C for 10 min, with a final extension at 72 °C for 2 min and a hold at 4 °C. The number of cycles was determined by a pilot PCR with 4–10 cycles of amplification, where we chose the minimum cycle number that produced a total DNA concentration of >50 ng/μL with no laddering. The resultant dsDNA was purified using a MinElute PCR Purification Kit (Qiagen) following the manufacturer’s protocol.

### Sample Pooling

In order to test multiple conditions on a single flow cell, we incorporated barcodes into the N_12_ region of the MIPs in order to enable multiplexing. Samples were prepared with varying input ratios of target molecules (**Table 2**). Post-RCA dsDNA products were pooled together before sequencing based on the measured concentration (HS dsDNA, Qubit) normalized by the median length reported by the 4200 Tape Station (Agilent).

### Nanopore sequencing & data analysis

The pooled library was prepared for sequencing via ONT’s Ligation Sequencing 1D Kit, and then sequenced on a R9.4 flow cell for 24 hrs according to the manufacturer’s protocol. Initial base-calling was done through ONT’s MinKNOW software. More sophisticated nanopore alignment tools were avoided as we wanted to test the worst-case scenario raw input, with as little data processing as possible required in between the nanopore run and HiFRe analysis. Custom MATLAB scripts (available at https://github.com/btotherad77/hifre) were developed to parse and analyze the raw nanopore reads. Individual repeats were identified as regions that mapped well to the FP sequence. Using the Smith-Waterman local alignment algorithm^32^, the mapping threshold was determined as the average alignment of random sequences to the initiator plus five standard deviations (*p* < 3 x 10^−7^). The identified individual repeats were collected and aligned to each other using a progressive multialignment algorithm (*multialign*, MATLAB). The consensus sequence was then determined base-wise with a winner-take-all strategy. The individual repeats and the consensus sequence were then compared back to the original template in order to assess the accuracy before and after alignment, respectively.

### Gel protocol

Gels were run using a PowerPac Basic (Bio-Rad) and imaged on a Molecular Imager Gel Doc XR+ (Bio-Rad). For amplified input DNA and post-RCA dsDNA, native gels were run with 5 μL per well of a 1:5:1 mixture of sample:water:BlueJuice loading dye (Thermo Fisher Scientific) in a 10% TBE, 1-mm, 12-well gel (Thermo Fisher Scientific) for 45 minutes at 150 V. For circularized ssDNA and post-RCA ssDNA, denaturing gels were run with 6 μL per well of a 1:1 mixture of sample and Gel Loading Buffer II with formamide (Ambion) in a 10% TBE-Urea, 1-mm, 12-well gel (Thermo Fisher Scientific) for 1 hr at 200 V. Sample-dye mixtures were initially denatured at 95 °C for 7 min.

## Acknowledgments

This work was financially supported by the Chan-Zuckerberg Biohub, and the Bill and Melinda Gates Foundation.

