## Supplemental Information for "High-Fidelity Nanopore Sequencing of Ultra-Short DNA Sequences"

Brandon D. Wilson[1], Michael Eisenstein[2,3], H. Tom Soh[2,3,4*]

**Affiliations:**

[1]Department of Chemical Engineering, Stanford University, Stanford, CA 94305, USA.

[2]Department of Electrical Engineering, Stanford University, Stanford, CA 94305, USA.

[3]Department of Radiology, Stanford University, Stanford, CA 94305, USA.

[4]Chan Zuckerberg Biohub, San Francisco, CA 94158, USA.

**Supplemental Calculation 1:** Determining the threshold for efficient circularization. We took care to design MIPs that are predicted to exhibit minimal secondary structure – especially in the hybridization regions – in order to ensure highly efficient circularization. Our threshold was based on the secondary structure thermodynamics that allowed for >90% conversion from linear to circular DNA over five temperature cycles (Equation 1):

$$\Delta G_{folding} = -RTln\left(\frac{1-f_{unfold}}{f_{unfold}}\right) \quad (1)$$

Since total conversion, $\varepsilon$, is related to the fraction of unfolded MIPs, $f_{unfold}$, and the cycle number, $n$, as shown in Equation 2:

$$\varepsilon = 1 - \left(1 - f_{unfold}\right)^n \quad (2)$$

we can calculate the minimum $\Delta G_{folding}$ of any individual motif (Equation 3):

$$\Delta G_{folding} \geq -RTln\left(\frac{(1-\varepsilon)^{1/n}}{1-(1-\varepsilon)^{1/n}}\right) \quad (3)$$

To achieve >90% circularization in 5 cycles, we must have $f_{unfold} > 0.37$. Therefore, $\Delta G_{folding}$ must be greater than $-0.33\,\frac{kcal}{mol}$ for any secondary structure motif involving one of the anchor sites. $\Delta G_{folding}$ was calculated on the *mfold* DNA folding form[33] at 37 °C and 10 mM $Mg^{2+}$. A secondary check was performed on NUPACK[32] to determine that $f_{unfold} > 0.37$ for all bases in both anchor sites.

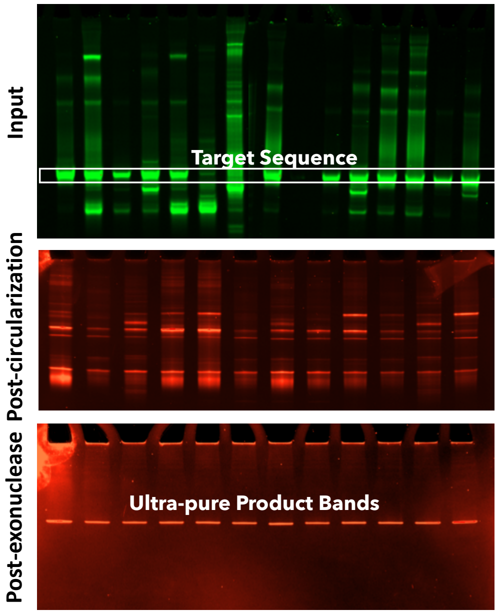


**Supplemental Figure 1:** MIPs can confer ultra-high specificity from a background of non-specific product. (top) PCR amplification can result in a large background of non-specific product, whereas (middle) circularization alone produces multiple bands. (bottom) Treatment with exonucleases I & III leaves only a single band of pure circular product. Top panel is a native TBE gel, whereas the other two panels are TBE-urea denaturing gels.

| Barcode | Sequence |
|---|---|
| b1 | 5'-TAGTCATCTCTA-3' |
| b2 | 5'-TACCAGGTCCTA-3' |
| b3 | 5'-TATCCATCCTTA-3' |
| b4 | 5'-TAAGCTCGCATA-3' |
| b5 | 5'-ATCCTCTCCTCA-3' |
| b6 | 5'-CGTCTACGATGC-3' |
| b7 | 5'-CACGAAGTGGAA-3' |
| b8 | 5'-GTTCCCTGTCCC-3' |
| b9 | 5'-ACGTTAAGGCCA-3' |
| b10 | 5'-TCCTCGTGAGGT-3' |
| b11 | 5'-CATCTAACCTAG-3' |
| b12 | 5'-TCGGAATTGGCT-3' |

**Supplemental Table 1:** List of barcodes used for demultiplexing. Barcodes are located in the $N_{12}$ region of the MIPs (**Error! Reference source not found.**)
